# Gene model for the ortholog of *Akt* in *Drosophila elegans*

**DOI:** 10.1101/2025.08.06.668967

**Authors:** Jhilam Dasgupta, Ashley Morgan, Anne E. Backlund, Juncheng Luo, Jefferson Waters, Jennifer A. Kennell

## Abstract

Gene model for the ortholog of *Akt kinase* (*Akt*) in the April 2013 (BCM-HGSC/Dele_2.0) Genome Assembly (GenBank Accession: GCA_000224195.2) of *Drosophila elegans*. This ortholog was characterized as part of a developing dataset to study the evolution of the Insulin/insulin-like growth factor signaling pathway (IIS) across the genus *Drosophila* using the Genomics Education Partnership gene annotation protocol for Course-based Undergraduate Research Experiences.

## Introduction

*This article reports a predicted gene model generated by undergraduate work using a structured gene model annotation protocol defined by the Genomics Education Partnership (GEP; thegep.org) for Course-based Undergraduate Research Experience (CURE). The following information in quotes may be repeated in other articles submitted by participants using the same GEP CURE protocol for annotating Drosophila species orthologs of Drosophila melanogaster genes in the insulin signaling pathway*.

“In this GEP CURE protocol students use web-based tools to manually annotate genes in non-model Drosophila species based on orthology to genes in the well-annotated model organism fruitfly Drosophila melanogaster. The GEP uses web-based tools to allow undergraduates to participate in course-based research by generating manual annotations of genes in non-model species (Rele et al., 2023). Computational-based gene predictions in any organism are often improved by careful manual annotation and curation, allowing for more accurate analyses of gene and genome evolution (Mudge and Harrow 2016; Tello-Ruiz et al., 2019). These models of orthologous genes across species, such as the one presented here, then provide a reliable basis for further evolutionary genomic analyses when made available to the scientific community.” (Myers et al., 2024).

“The particular gene ortholog described here was characterized as part of a developing dataset to study the evolution of the Insulin/insulin-like growth factor signaling pathway (IIS) across the genus *Drosophila*. The Insulin/insulin-like growth factor signaling pathway (IIS) is a highly conserved signaling pathway in animals and is central to mediating organismal responses to nutrients (Hietakangas and Cohen 2009; Grewal 2009).” (Myers et al., 2024).

“*Akt kinase* (*Akt* also known as *Akt1, Protein Kinase B, PKB;* FBgn0010379) regulates stress response, aging, and cell growth and survival in *Drosophila* (Stavely et al., 1998; Verdu et al., 1999). It is involved in signal transduction pathways in physiological and neurological pathways in *Drosophila* (Guo and Zhong 2006). It encodes a core serine-threonine kinase (Bellacosa et al. 1991) component of the Insulin-like growth factor pathway that functions downstream of, and following its activation by the *Pi3K92E* product in *Drosophila* (Andjelkovic et al., 1995). It is activated by phosphatidylinositol binding and phosphorylation (Potter et al., 2002).” (Morgan et al., 2022).

We propose a gene model for the *D. elegans* (NCBI:txid30023) ortholog of the *D. melanogaster Akt kinase* (*Akt*) gene. The genomic region of the ortholog corresponds to the uncharacterized protein LOC108140670 (RefSeq accession XP_017119102.1) in the April 2013 (BCM-HGSC/Dele_2.0) Genome Assembly of *Drosophila elegans* (GenBank Accession: GCA_000224195.2). This model is based on RNA-Seq data from *D. elegans* (PRJNA63469; modENCODE Consortium et al., 2011; Chen et al. 2014) and *Akt* in *D. melanogaster* using FlyBase release FB2022_04 (GCA_000001215.4; Gramates et al., 2022; Jenkins et al., 2022; Larkin et al., 2021).

### Synteny

The reference gene, *Akt*, occurs on chromosome 3R in *D. melanogaster* and is flanked upstream by *mini spindles* (*msps*), *CG10185*, and *Myosin heavy chain-like* (*Mhcl*) which nests *CG32855* and *sex-specific enzyme 2* (*sxe2*) and downstream by *Stubble* (*Sb*) and *MICOS subunit 26/27* (*Mic26-27*). The *tblastn* search of *D. melanogaster* Akt-PC (query) against the *D. elegans* (GenBank Accession: GCA_000224195.2) Genome Assembly (database) placed the putative ortholog of *Akt* within scaffold scf7180000491080 (KB458458.1) at locus LOC108140670 (XP_017119102.1)— with an E-value of 3e-149 and a percent identity of 73.50%. Furthermore, the putative ortholog is flanked upstream by LOC108140553 (XP_017118899.1), LOC108140555 (XP_017118903.1), and LOC108140512 (XP_017118792.1), which nests LOC108140512 (XP_017118795.1) and LOC108140514 (XP_017118803.1). These correspond to *msps, CG10185*, and *Mhcl* which nests *CG32855* and *sxe2* in *D. melanogaster* (E-value: 0.0, 0.0, 0.0, 0.0, and 0.0; identity: 93.88%, 97.69%, 96.64%, 83.39%, and 84.75%, respectively, as determined by *blastp*; Figure 1A, Altschul et al., 1990). The putative ortholog of *Akt* is flanked downstream by LOC108140647 (XP_017119062.1) and LOC108140648 (XP_017119063.1), which correspond to *Sb* and *Mic26-27* in *D. melanogaster* (E-value: 0.0 and 4e-157; identity: 84.05% and 92.14%, respectively, as determined by *blastp*). The putative ortholog assignment for *Akt* in *D. elegans* is supported by the following evidence: The genes surrounding the *Akt* ortholog are orthologous to the genes at the same locus in *D. melanogaster* and synteny is completely conserved, supported by results generated by *blastp*; we conclude that LOC108140670 is the correct ortholog of *Akt* in *D. elegans* (Figure 1A).

**Figure 1:**
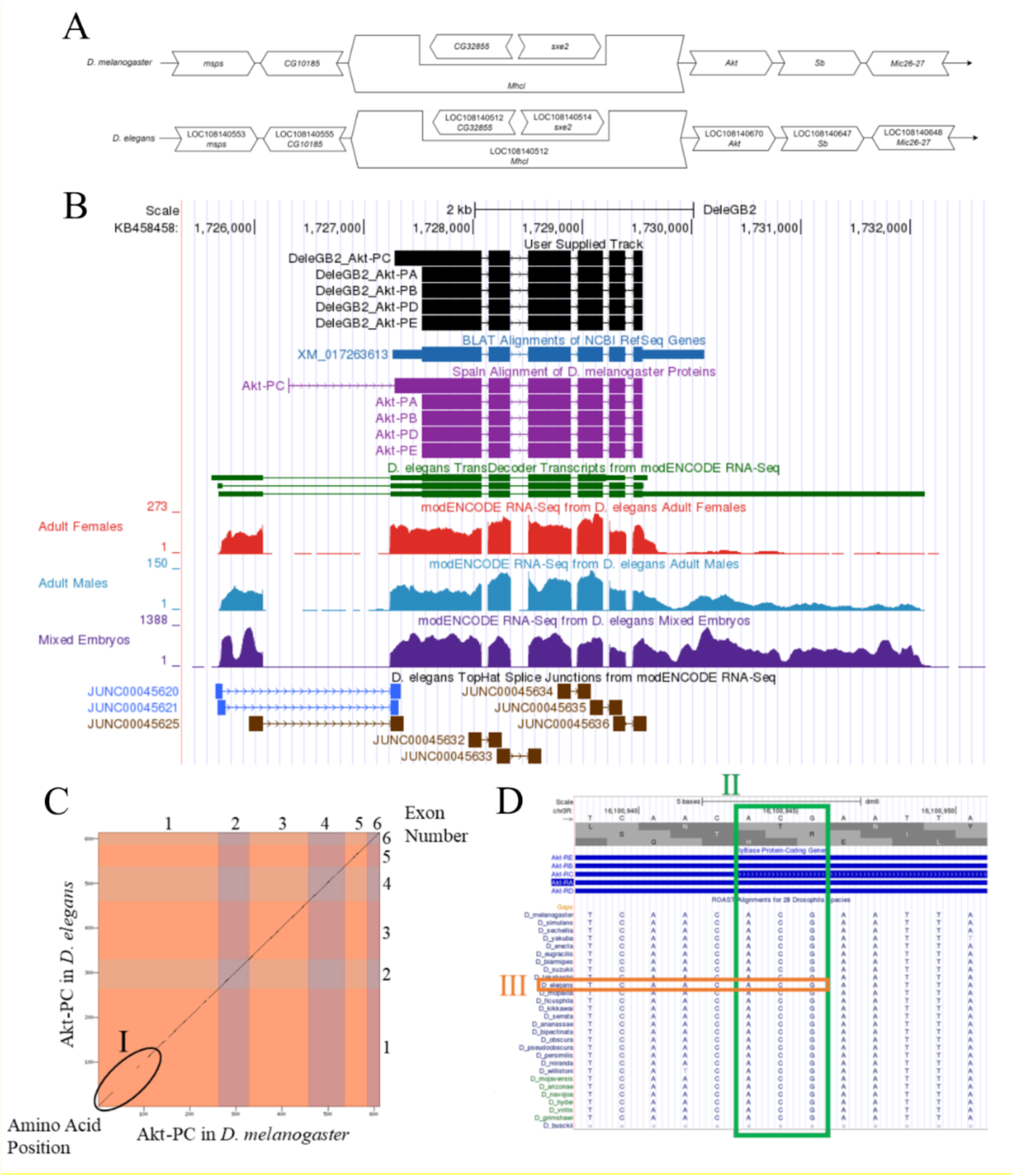
Akt gene model comparison between *Drosophila elegans* and *Drosophila melanogaster* orthologs. **(A) Synteny comparison of the genomic neighborhoods for *Akt* in *Drosophila melanogaster* and *D. elegans***. Thin underlying arrows indicate the DNA strand within which the reference gene–*Akt*–is located in *D. melanogaster* (top) and *D. elegans* (bottom). The thin arrows pointing to the right indicate that *Akt* is on the positive (+) strand in *D. elegans* and *D. melanogaster*. The wide gene arrows pointing in the same direction as *Akt* are on the same strand relative to the thin underlying arrows, while wide gene arrows pointing in the opposite direction of *Akt* are on the opposite strand relative to the thin underlying arrows. White gene arrows in *D. elegans* indicate orthology to the corresponding gene in *D. melanogaster*. Gene symbols given in the *D. elegans* gene arrows indicate the orthologous gene in *D. melanogaster*, while the locus identifiers are specific to *D. elegans*. **(B) Gene Model in GEP UCSC Track Data Hub (Raney et al**., **2014)**. The coding-regions of *Akt* in *D. elegans* are displayed in the User Supplied Track (black); coding exons are depicted by thick rectangles and introns by thin lines with arrows indicating the direction of transcription. Subsequent evidence tracks include BLAT Alignments of NCBI RefSeq Genes (dark blue, alignment of Ref-Seq genes for *D. elegans*), Spaln of D. melanogaster Proteins (light purple, alignment of Ref-Seq proteins from *D. melanogaster*), Transcripts and Coding Regions Predicted by TransDecoder (dark green), RNA-Seq from Adult Females, Adult Males, and Mixed Embryos (red, light blue, dark purple, respectively; alignment of Illumina RNA-Seq reads from *D. elegans*), and Splice Junctions Predicted by regtools using *D. elegans* RNA-Seq (PRJNA63469). Splice junctions shown have a read-depth of 10-49 and 500-999, supporting reads in blue and brown, respectively. **(C) Dot Plot of Akt-PC in *D. melanogaster* (*x*-axis) vs. the orthologous peptide in *D. elegans* (*y*-axis)**. Amino acid number is indicated along the left and bottom; coding-exon number is indicated along the top and right, and exons are also highlighted with alternating colors. Region I encircled in black indicates lack of sequence similarity between the two sequences. **(D) ROAST Alignments Conservation Track Displaying Non-Canonical (ACG) Start Codon within the UCSC Genome Browser**. The evidence tracks displayed are FlyBase Protein-Coding Genes (blue) and ROAST Alignment for 28 Drosophila Species. Green box II outlines the location of the non-canonical start codon in *D. melanogaster* as well as the sequences of other species in this location. Orange box III highlights the nucleotides in *D. elegans* at this location.

### Protein Model

*Akt* in *D. elegans* has five protein-coding isoforms (Akt-PA, Akt-PB, Akt-PC, Akt-PD, Akt-PE) which all contain six protein-coding exons. Four of these isoforms have identical protein-coding exons (Akt-PA, Akt-PB, Akt-PD, Akt-PE; Figure 1B), and isoform (Akt-PC) differs in the length of the first exon (Figure 1B). Relative to the ortholog in *D. melanogaster*, the coding-exon number is conserved. The sequence of Akt-PC in *D. elegans* has 93.95% identity (E-value: 0.0) with the protein-coding isoform Akt-PC in *D. melanogaster*, as determined by *blastp* (Figure 1C). The non-canonical start codon (ACG) which exists in the Akt-PC isoform in *D. melanogaster* was also found in the Akt-PC isoform in *D. elegans*. This gene model can be seen within the target genome at this TrackHub.

#### Special characteristics of the protein model

Akt-PC isoform has a non-canonical ACG start codon in *D. melanogaster*. This start codon is maintained throughout 28 *Drosophila* species (Figure 1D). The level of conservation of the ACG start codon across all 28 *Drosophila* species, and the general region surrounding the non-canonical codon, leads us to conclude that the Akt-PC isoform also exists in *D. elegans*.

## Methods

“Detailed methods including algorithms, database versions, and citations for the complete annotation process can be found in Rele et al. (2023). Briefly, students use the GEP instance of the UCSC Genome Browser v.435 (https://gander.wustl.edu; Kent WJ et al., 2002; Navarro Gonzalez et al., 2021) to examine the genomic neighborhood of their reference IIS gene in the *D. melanogaster* genome assembly (Aug. 2014; BDGP Release 6 + ISO1 MT/dm6). Students then retrieve the protein sequence for the *D. melanogaster* reference gene for a given isoform and run it using *tblastn* against their target *Drosophila* species genome assembly on the NCBI BLAST server (https://blast.ncbi.nlm.nih.gov/Blast.cgi; Altschul et al., 1990) to identify potential orthologs. To validate the potential ortholog, students compare the local genomic neighborhood of their potential ortholog with the genomic neighborhood of their reference gene in *D. melanogaster*. This local synteny analysis includes at minimum the two upstream and downstream genes relative to their putative ortholog. They also explore other sets of genomic evidence using multiple alignment tracks in the Genome Browser, including BLAT alignments of RefSeq Genes, Spaln alignment of *D. melanogaster* proteins, multiple gene prediction tracks (e.g., GeMoMa, Geneid, Augustus), and modENCODE RNA-Seq from the target species. Detailed explanation of how these lines of genomic evidenced are leveraged by students in gene model development are described in Rele et al. (2023). Genomic structure information (e.g., CDSs, intron-exon number and boundaries, number of isoforms) for the *D. melanogaster* reference gene is retrieved through the Gene Record Finder (https://gander.wustl.edu/~wilson/dmelgenerecord/index.html; Rele et al., 2023).

Approximate splice sites within the target gene are determined using *tblastn* using the CDSs from the *D. melanogaste*r reference gene. Coordinates of CDSs are then refined by examining aligned modENCODE RNA-Seq data, and by applying paradigms of molecular biology such as identifying canonical splice site sequences and ensuring the maintenance of an open reading frame across hypothesized splice sites. Students then confirm the biological validity of their target gene model using the Gene Model Checker (https://gander.wustl.edu/~wilson/dmelgenerecord/index.html; Rele et al., 2023), which compares the structure and translated sequence from their hypothesized target gene model against the *D. melanogaster* reference gene model. At least two independent models for a gene are generated by students under mentorship of their faculty course instructors. Those models are then reconciled by a third independent researcher mentored by the project leaders to produce the final model. Note: comparison of 5’ and 3’ UTR sequence information is not included in this GEP CURE protocol.” (Gruys et al., 2025)

## Supporting information

Gene model data files

## Supplemental Files

1. Zip file containing a FASTA, PEP, GFF files for the gene model
2. Figure 1 in high resolution

### Metadata

Bioinformatics, Genomics, *Drosophila*, Genotype Data, New Finding

## Acknowledgements

We would like to thank Wilson Leung for developing and maintaining the technological infrastructure that was used to create this gene model and Laura K. Reed for overseeing the project. Thank you to FlyBase for providing the definitive database for *Drosophila melanogaster* gene models. Further, we would like to thank the editors and developers at the journal *microPublication: Biology* for assistance in developing the template for these single gene ortholog publications.

